# Multimodal interactions in Stomoxys navigation reveals synergy between olfaction and vision

**DOI:** 10.1101/2024.01.11.575173

**Authors:** Merid N Getahun, Steve Baleba, John Ngiela, Peter Ahuya, Daniel Masiga

## Abstract

Stomoxys flies exhibit an attraction towards objects that offer no rewards, such as traps and targets devoid of blood or nectar incentives. This behavior provides an opportunity to develop effective tools for vector control and monitoring. However, for these systems to be sustainable and eco-friendly, the visual cues used must be selective in attracting the target vector(s). In this study, we modified the existing blue Vavoua trap, originally designed to attract biting flies, to create a deceptive host attraction system specifically biased towards attracting Stomoxys. Our research reveals that Stomoxys flies are attracted to various colors, with red proving to be the most attractive and selective color for Stomoxys compared to other colors tested. Interestingly, our investigation on cattle-Stomoxys interaction demonstrates that Stomoxys flies do not have preference for a specific livestock fur color phenotype, despite variation in spectrum. To create a realistic sensory impression of the trap in the Stomoxys nervous system, we incorporated olfactory cues from livestock host odors that significantly increased trap catches. The optimized novel nanopolymer bead dispenser capable of effectively releasing the attractive odor, carvone + p-cresol, with strong plume strands, longevity. Overall, red trap baited with nano polymer beads dispenser is environmentally preferred.

## Introduction

When insect vectors make use of multimodal signals such as host scent, color, morphology, auditory, gustatory, mechanosensory signals at different time in space help them to minimize the mistake and make almost perfect decision in locating their blood meal source, nectar, mate partner [1–5]. However, due to the nature of the signals variation in space and time insects use some of the signal(s) at different time in space, for instance at far distance with a lot of visual background signals such as in a forest or bushy environment olfactory cue plays a significant role as there are barrier to resolve visual cues [6]. Such behavior, i.e., the use of individual signal or minimum cue(s), reduction approach to represent a given host make insect vulnerable for deception. Insects make use of their visual signal to perform various behaviors, including flight control, object tracking for host or nectar-finding and have preference for certain bands from the visible spectrum inputs for their ecological interaction, including to get blood meal source and thus for disease transmission [7] [8–12]. When we compare the natural deception system by those plants to be pollinated by insects without rewarding nectar the plants evolved to generate a perfect sensory impression in terms of smell, shape and even heat [13] [14] of a desirable host in the insect nervous system. However, in biting flies such as Stomoxys, tabanids and tsetse flies using a simple target and trap of blue color that does not look or smell like a cow, can easily catch a good number of hungry biting flies [10,11,15–19].

However, supporting the multimodal signals principle the deception can be significantly enhanced by adding additional inputs, for instance addition of host scent alone exhibited by the significant increase in trap catch in tsetse flies and Stomoxys [6,19–21]. However, there is variation between vectors in deception, for instance kissing bugs prefer visual objects only when baited with odors [22]. *Aedes aegypti* is not attracted to black objects in the absence of CO_2_, but after encountering a CO_2_ plume, they become highly attracted to such objects [23] demonstrating the importance of various signals integration and variation between insect for visual object attraction. Historically, the design of traps for biting flies has primarily focused on maximizing trap catch, with less emphasis placed on selectivity. For instance, [21,24–26], demonstrated high diversity of insects caught in biting flies trap, such as in Vavoua, Nzi traps including non-target insects. This highlights the need for improvement in making biting flies traps more selective. The potential role of various livestock fur colors in influencing stable fly-livestock interactions, and the development of odor dispensers to enhance the sensory impression of the trap to Stomoxys, have received limited attention. In this study, we address these gaps by modifying the Vavoua trap’s blue color [1] to red, demonstrating its selectivity towards Stomoxys flies without compromising its effectiveness in capturing Stomoxys. Furthermore, we have optimized a novel nanopolymer bead dispenser to release livestock host odors, thereby increasing the trap’s efficiency.

### Materials Fabric Colors

Indigenous African livestock exhibit a wide range of genotypes[27], which is reflected in their diverse fur color phenotypes that may impact their interactions with biting flies (Fig. 1A). Additionally, Stomoxys flies feed on various nectar sources[28–30], which themselves display a diverse array of flower colors (Fig. 1D-F). To investigate fabric colors that could potentially be more selective for attracting Stomoxys we conducted a field study using eight colors in polyester-cotton fabrics bought from local market in Nairobi, Kenya. Some of these colors were chosen to resemble plant leaves, flowers, or animal skin color, while blue was used as a positive control (Fig. 1E).

**Fig. 1.**
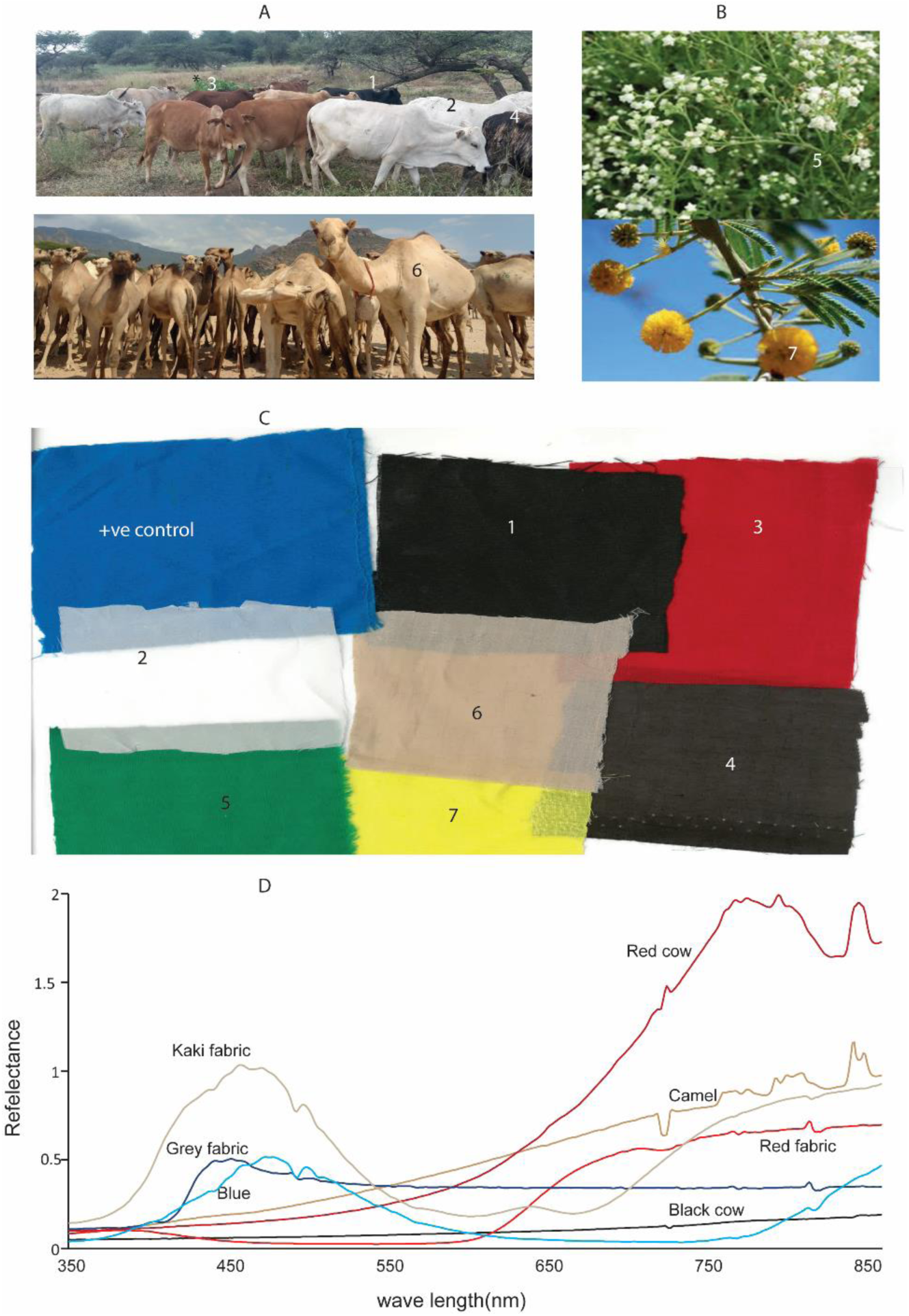
Livestock fur Skin color and nectar source of plants and the corresponding fabric color to represented livestock and plants parts. (A) Photo showing the various color phenotypes of livestock, blood meal source of Stomoxys (B) nectar source (Photo icipe/MNG). (C) The various fabric colors used for behavioral evaluation. (D) spectrophotometer measurement from selected livestock fur color and selected used fabric. The number matching shows how we represented animals’ fur and plants color with fabric color.

### Chemicals

We used pure (R)-(−)-carvone, an odor known to attract gravid stomoxys flies [31] and p-cresol livestock derived semio-chemicals that attract blood seeking Stomoxys [21,32]. The chemicals were obtained from Sigma Aldrich Germany, R-(−)-carvone (98%) and p-cresol (98%) purity. These two odors were chosen to formulate a blend with 1:1 ratio for potential synergism.

### Dispensers

The present study aimed to examine the suitability of paraffin wax and nanopolymer beads as carrier materials for attractants in field applications.

### Wax dispenser making

Odorless Paraffin wax was obtained from (Nairobi Pharmaceuticals) 15 ml of the wax was heated at 60 °C to melt, then 800 µl of the blend of 1:1 ratio of p-cresol and carvone was dissolved in 15 ml wax and mixed for 30 second and the liquid was poured into a mold and allowed to solidify to make wax dispenser. The loaded dispensers were left under field conditions and taken to the lab for inherent release measurement. For odor trapping a general purpose 65μm PDMS/DVB (polydimethyl siloxane/divinylbezene, Supelco, Bellefonte, PA, USA, SPME fibers were used[33]. The SPME fibers for adsorbing the odors were placed directly above the wax. The inherent release characteristics of the wax to the blend was evaluated from day zero for six days.

### Nanopolymer beads dispenser

Nanopolymer beads product number 20009316 obtained from Celanese EVA Performance Polymers Inc. Canada. Equal amount of 800 µl of the blend was impregnated in 4gm beads for 24 hours under hood with frequent shaking for some time, then the beads were placed in a12 cm long circular tygon tube with 0.635cm internal diameter, 0.953cm external diameter and 0.159cm wall thickness; (Cole Parmer International). We used SPME to mimic insect antenna to measure the amount of odors plume strand flux that the SPME and therefore an encountering insect’s antenna would encounter when these dispensers dispensed the given odors the same as[34]. We measure the plume strand on different days, from day 0 daily for six days the same as above by placing the SPME directly above the impregnated beads.

### Electrophysiology

We used *S.calcitrans* as representative to measure the response of the olfactory sensory neurons to the compounds using Electroantennography (EAG) [35] the techniques that measures the sum total of electrical potentials generated by activated Olfactory Sensory Neurons (OSNs) on the insect antenna. We used 10 impregnated beads placed in glass pipet to stimulate the OSNs, with the following treatments: unimpregnated beads used as a control, p-cresol impregnated beads, carvone impregnated beads and blend impregnated beads, we also tested blend stayed under field condition for 7 days to see if that affect the response as compared to newly formulated blend. We measured the olfactory receptor potentials from the whole insect inserted in 1000ul pipet tip and the head is pushed out to get access to the antenna. Glass capillary microelectrodes with silver electrode, filled with ringer solution, 6.4 mM KCl, 20 mM KH 2 PO 4, 12 mM MgCl 2, 1 mM CaCl 2, 9.6 mM KOH, 354 mM glucose, 12 mM, NaCl, pH 6.5 [36] inserted in the eye for grounding and the other recording electrode at the tip of the antenna, a slight cut was made at the tip of the antenna to establish electric connection and the recording electrode was connected to an 10X amplifier and a recording instrument. We stimulated the antenna with 500ms pulse duration.

### Field trapping

To evaluate the attractivity of various fabric color we conducted 8 x 8 Latin square design experiment at Mpala Ranch located in Laikipia County Central Kenya field site, which was previously described[31]. We used a 20 by 20 cm small square target covered with Rentokil sticky material on both sides and hung 30 cm above the ground. Initially the color was assigned randomly and everyday shifted to the new position to avoid any position effect. Similarly, we used a 4×4 Latin square design experiment to evaluate the efficacy of the modified traps and dispensers at Ngurunit Northern Kenya, Isiolo, Shimba Hill Coastal Kenya and Gatundu around Nairobi area. For cattle-Stomoxys interaction to see if Stomoxys flies have preference for certain livestock fur color we counted number of Stomoxys flies on various fur-colored cattle from two sites (Isiolo and Nguruman), these are two sites among our sites with more cattle populations. Flies caught in the traps were identified morphologically according to [37].

### Livestock reflectance measurement

The measurements of the livestock fur spectrum were obtained by positioning the measuring device 20-30cm above the animals’ backs. This was done in the morning 9-11am under clear sky conditions, after the animals had been resting on the ground. We employed the in-situ FieldSpec® Handheld 2™ analytical spectral instrument (ASD-USA) the same as [38]. The spectroradiometer was configured to internally and automatically gathered and compute an average of 20 spectral measurements for every measurement of the sample spectrum. We conducted measurements from three to five animals with identical fur color, following optimization and calibration of the measured radiance. This was achieved by utilizing a Spectralon white reference with about 100% reflectance. For fabric, we utilized Spectroradiometer RS-8800 USA is a handheld non-imaging high spectral resolution/high sensitivity system and captures a full spectral range (350-2500 nm).

In orderto obtain the measurement, we obtained a small square cloth measuring 10 × 10 cm and positioned it on the table and measurement was done the same as the livestock fur. Prior to each fabric measurement, we standardized the reading by comparing it to the measurement taken against a white backdrop.

### Data analysis

To compare the number of Stomoxys spp. caught by the different colored target, we ran the Kruskall Walliss nonparametric test followed by the Dunn post-hoc statistical tests, as data were not normally distributed (Shapiro-Wilk test: p<0.05) and variance was not homogeneous (Levene test: p<0.05). For color selectivity, all insects’ orders caught except house flies and stomoxys were pooled together and considered as non-target and then compared against stomoxys catch using independent t-test or Mann Whitney test depending on the data normality. We applied one way ANOVA to compare more than two independent treatments, and we used PRISM 9.04 to analyze the data. All statistical results were considered significant at p < 0.05.

## Results

### Skin color of livestock and plants nectar source varies in their wavelength

Stomoxys feeds both on blood and nectars [28,30,39]. For example, cattle demonstrate various phenotypes in their skin color, from black, brown, white, various reddish (Fig 1A). However, *Camelus dromedarius* fur color is dominated by camel color which is represented by kaki fabric color (Fig.1B), the color variation demonstrated in their spectrum (Fig.1G). Black and dark brown colored cattle have low reflectance across wavelength, but other colored livestock starts increasing around wavelength of 600nm. Things to note in livestock spectrum, there is no spectrum shape like that of fabric, low at UV, rise in the visible spectrum (400-700 depending on color) and then fall at infrared zone, the spectrum is straight line increasing in reflectance as we move from UV, visible and infrared light spectrum, that means the reflectance steadily increases from 300 to 700 nm and shows no spectral peak data (Fig.1D, Supplementary table 1). Similarly, plants leaves and flowers source of nectar varies in their color (Fig.1B), from green, red, yellow, to white they have low in UV and have reflectance between 500-600 and high infrared reflectance (data not shown).

### Stomoxys flies are attracted to various colored sticky targets

In previous studies blue Vavoua trap developed by[16] was found to be effective for biting flies sampling, however, nontarget insects were trapped[21,24]. We ask if we test more colors, resembling their host, blood meal and nectar source color may minimize the catch of nontarget insects. We found there is a significant difference between colors in attracting stomoxys, Kruskal Wallis test, P<0.001. With pair comparison sticky targets with red color followed by kaki, blue, and white/grey were more attractive to Stomoxys spp. as compared to the other tested colors (Fig. 2A). Yellow and green were found to be less attractive. Furthermore, based on the analysis of each color to nontarget insects identified at the order level (Hymenoptera, Lepidoptera, Coleoptera, and Orthoptera), red color was more selective to Stomoxys as compared to other tested colors in attracting nontarget insects, Mann Whitney test, P=0.016 (Fig.2H). While blue, Kaki and white/grey they were equally attractive to other non-target insects P > 0.05 independent t-test (Fig 2B-G and I, Supplementary table 2). Independent of the sticky target colours, we significantly caught more dipteran insects as compared to other insect orders.

**Figure 2.**
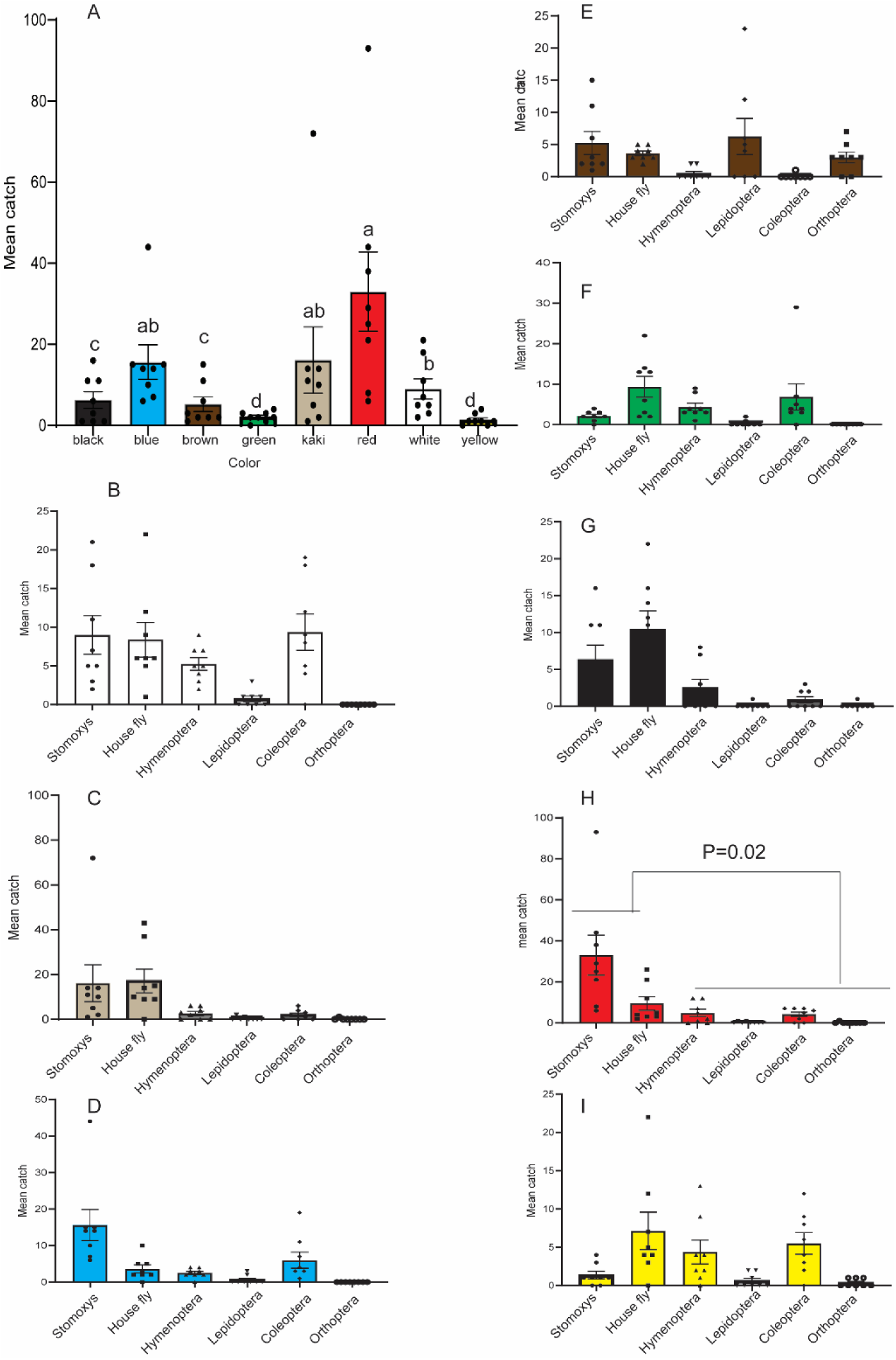
The attractivity of various colored small sticky target to Stomoxys and other insects. (A) Graph depicting the variation of Stomoxys flies catches across the different colored targets. (B-I) graphs illustrate the attractiveness of different colors to diverse insect groups. The graph shows the attractivity of the given color to various insect group. The error bar shows standard error of the mean. In Fig. 2A Bars with different letters are significantly different from each other based on the Kruskall analysis followed by Dunn post-hoc test. 2H, shows significant difference between Stomoxys and pooled non-target insects catch.

### Stomoxys and cattle visual interaction

We then asked if Stomoxys spp. have any color preference for various cattle phenotype, in this case livestock fur color (Fig1A). We have counted the number of Stomoxys from five various available colors in a given herd, at two sites Isiolo and Nguruman while cattle are inside their boma’s assisted by photo and video. We found that Stomoxys aggregate and feed on lower legs and around head, while feeding, they generally position themselves facing up-ward (Fig.3A inset) and tend to feed on all livestock color no statistical difference was observed at Isiolo (Fig.3A), ANOVA, F=0.7125, P=0.59. Per cow up to 115 Stomoxys flies were observed feeding at a given time. However, there is a slight variation at Nguruman sites (Fig. 3B), ANOVA F, 3.37, P=0.02.

**Figure 3.**
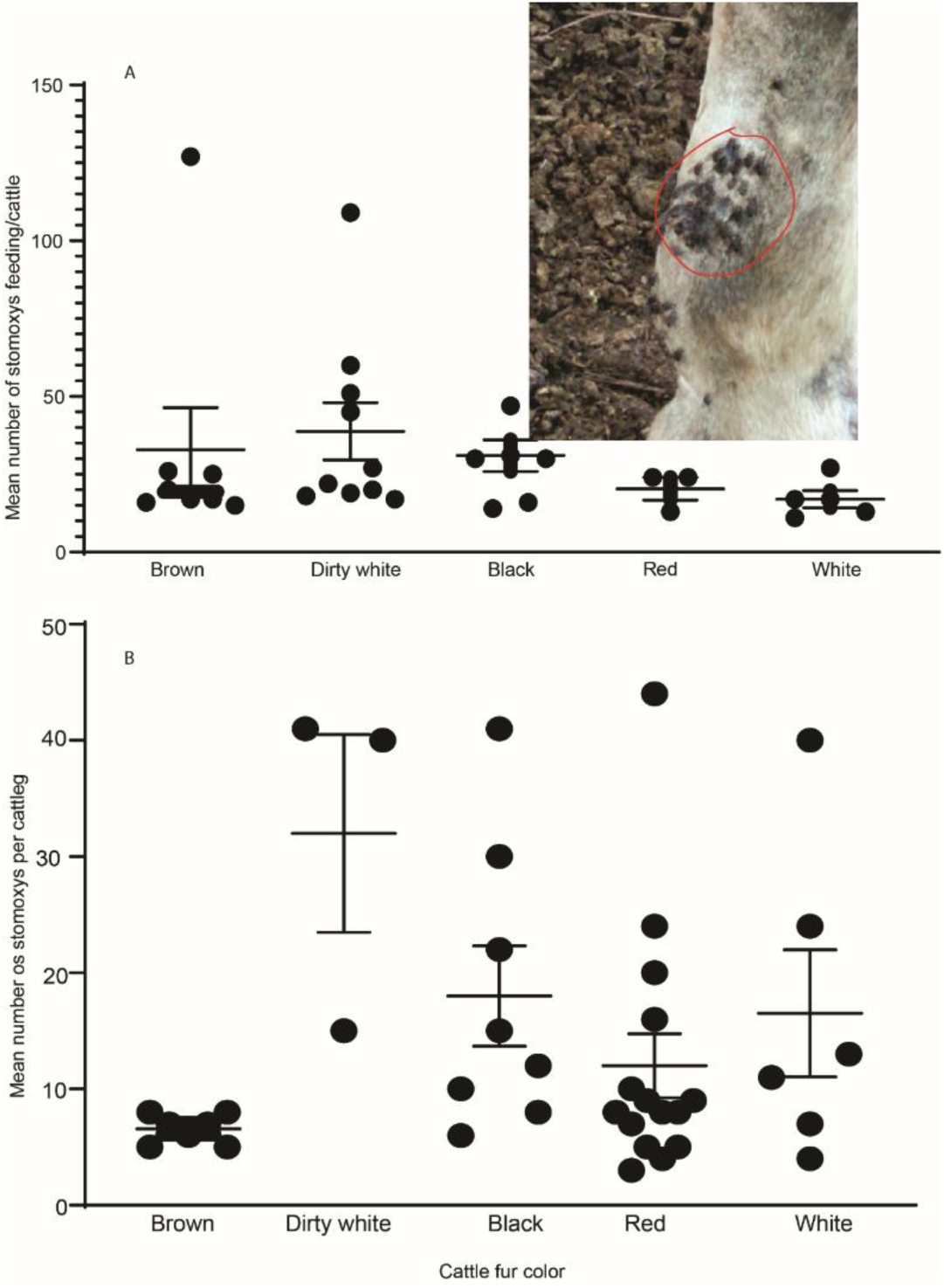
cattle-Stomoxys interaction (A) at Isiolo site and (B) at Nguruman site

### Making Vavoua trap stomoxys flies specific

We then replaced the Vestergaard blue color in Vavoua trap (zero fly) with red cotton-polyster fabric color from locally available fabric, the trap was made locally (Fig.4A) and tested the attractivity of the red vs Vestergaard blue Vavoua trap (zero fly) under field condition. We were able to replicate the result of the tiny target, as both red and blue were equally attractive to both Stomoxys spp. and house flies (Fig.4B-C), no significant difference between red and blue colored traps in catching Stomoxys spp. and house fly, independent t-test, P > 0.05 (Fig.4B). Like the target fabric, Vavoua trap designed with red color is more selective to Stomoxys flies as compared to the blue Vavoua trap (Fig.4B). More nontargets insects such as Hymenoptera, Lepidoptera and coleoptera were caught in blue Vavoua trap as compared to the red Vavoua trap, independent t-test, P<0.05. Similarly, at Isiolo site both traps attracted equal number of Stomoxys, t=0.477, p=0.6, house flies (t=1.09, P=0.3) similarly red attracted only 0.76x of the non-target insects as compared to blue colored trap. However, we acknowledge the number of non-target insects was low (Fig.4C). At Gatundu, we found both colors competitive, both traps caught specifically Stomoxys flies (Fig 4D). In all sites we encountered mainly three species of Stomoxys, *S.calcitrans*, Linnaeus *S.niger*, Linnaeus and *S. buati*, with varying proportion, but the first two are the most dominant species.

**Fig. 4.**
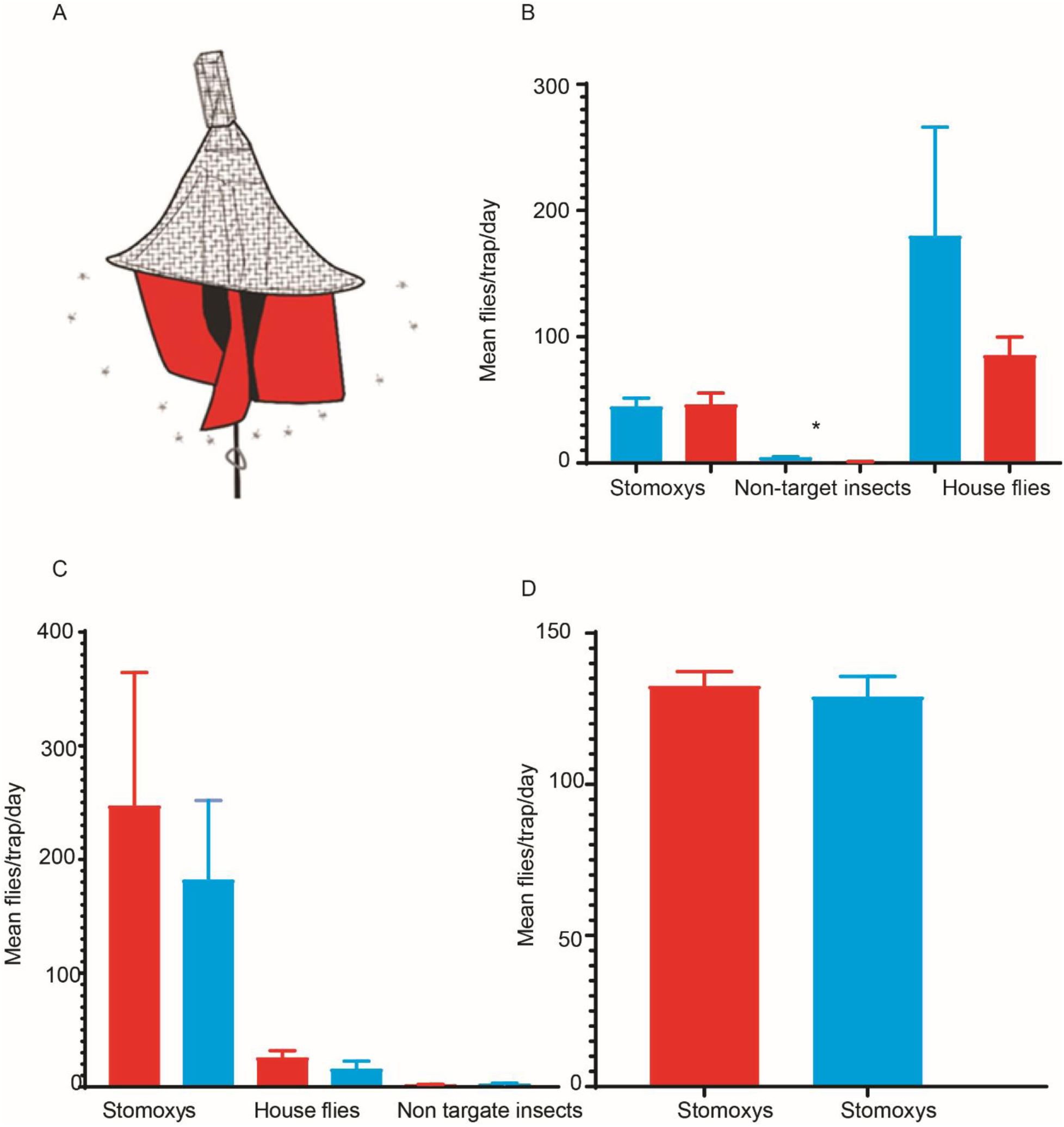
Attractivity of unbaited red and blue monoconcial traps to various insects at three different sites. (A) sketch of the modified monoconcial trap (B) Mean trap catch of Stomoxys flies, house fly and nontarget insects between red and blue traps under field condition. * Depicts a significant difference in catch of non-target insets, independent t-test, p< 0.05 at Ngurunit site. (C). Catch of various insects at Isiolo site. (D) Stomoxys catch from Gatundo, Nairobi area.

### Nanopolymer beads created a strong strands and controlled release of semio-chemicals

The two dispensers (Fig. 5A) were compared for the odor releases and attractivity under field condition. The nanobead formulation produced strong odor strand as compared to the wax formulation, see area under the curve of GC-MS chromatograph (Fig. 5B-D). Furthermore, these two dispensers also vary in their odor release, the odor release was odor and dispenser specific. For instance, the wax formulation lost 50% of p-cresol after 96 Hr, while nanobead lost only 20% of p-cresol. While carvone loss was ∼ 67% for wax but only ∼13% for the nanobead formulation. Based on GC-MS peak intensity wax released both compounds with equal ratio at day 0, but in beads p-cresol was low as compared to carvone (Fig. 5B). Based on the odors - carrier interaction nanobeads carrier created strong strand and controlled release that is reflected in behavioural response efficacy (Fig 5E). Seven-day field trapping experiment shows the nanobead formulation was more attractive as compared to the wax formulation across days (Fig.5E) the number of flies was fluctuating between days. This demonstrates the nano polymer bead delivered more constant release-rate with strong strand for over longer period that has improved behavioural response and significantly enhanced trap attractivity. In addition to the high release rate the wax dispenser at day 7 melted, so not suitable for arid and semi-arid areas or in hot environment. Thus, we used the nanopolymer beads for further experiments. The amount of the two odors after 15 days in the field was reduced only by 50% for p-cresol and 35 % to that of carvone when nanobeads utilized as dispenser.

**Fig 5.**
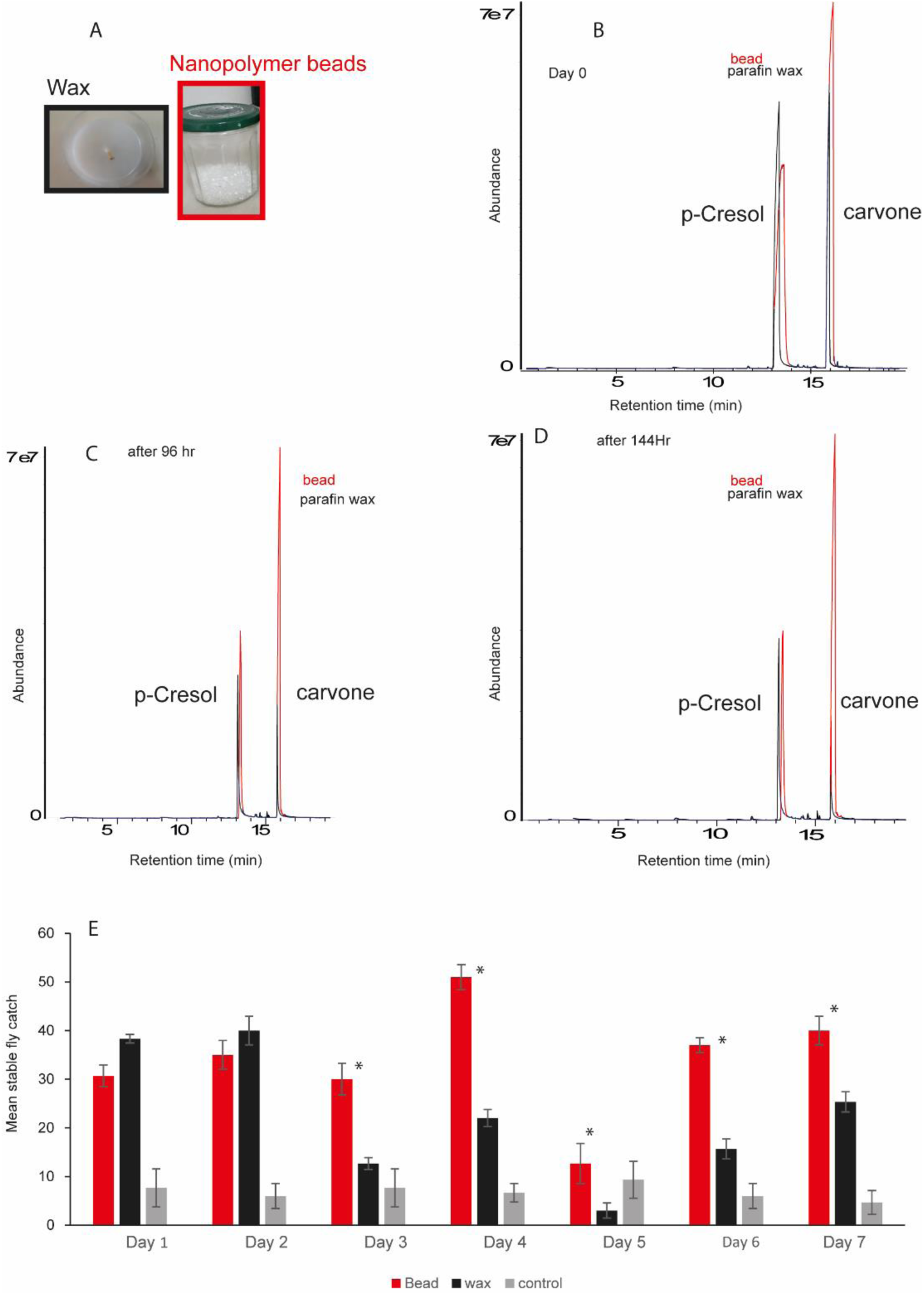
Attractant release and attractivity depends on dispenser type. (A) The two dispensers used (B) Blend odor strand from the two dispensers as it is released at day 0 (before placed under field condition). (C) Blend odor strand from the two dispensers after day 4 under field conditions (D) the same on day 6 (E). Mean Stomoxys catch between the three treatments. * Depicts a significant difference between nanobeads and wax dispenser, independent t-test. The trap catch was done at icipe campus Nairobi.

### Synergy of blend formulation

We formulated a blend of carvone and p-cresol, to target both blood meal searching as well as gravid females for maximum impact, the blend constituted with 1:1 ratio of each and impregnated in nanobeads. First, we asked if blend has any synergism effect on trap catch. We observed that the exposure of the blend for seven days did not affect the olfactory sensory neurons response, the mean mV of the blend at day zero and used blend at day seven was the same, t-test, t=0.2566, df=18, p=0.8 (Fig.6A-C, Supplementary Table 3). Under field condition we found that the blend formulation attracted more Stomoxys as compared to individual components, F=7.486, P=0.012, however, there was no difference between the two individual compounds (Fig.6D). Unlike the behavioral response we did not see olfactory sensory neurons response synergism due to blend, at the antennal level response (Fig.6A-B). However, we observed the response duration or recovery rate is shorter in blends (new and used) as compared to carvone, Kruskal-Wallis test, P<0.0001(Fig.6C), but no difference with p-cresol. Impregnating the odours in nanobeads exhibits enhanced behavioral efficacy and maintains long-lasting performance in field conditions, even when using reduced odour loading rates per dispenser.

**Fig. 6.**
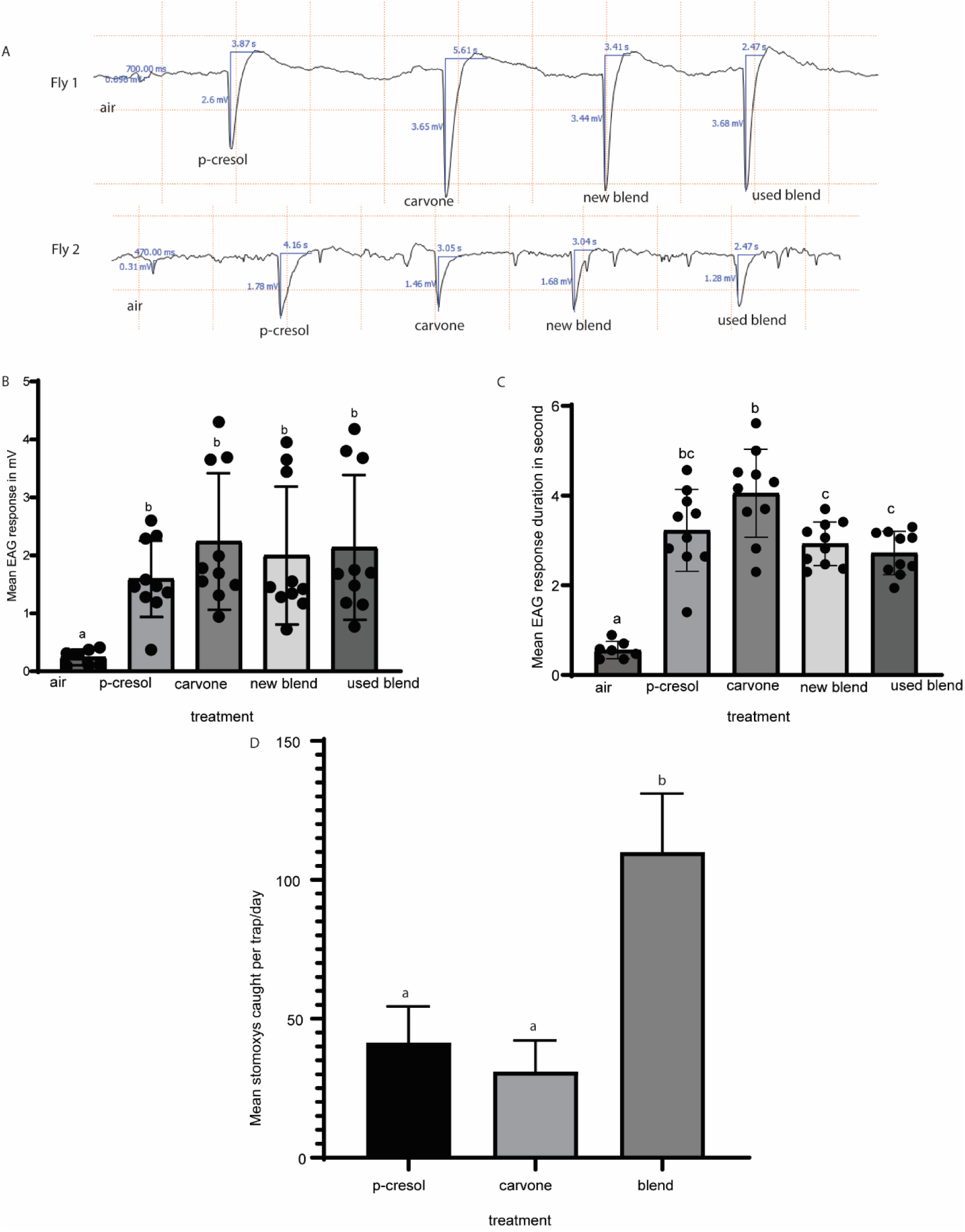
Electrophysiological and behavioral response of Stomoxys to single and blend formulation. (A) Representative antennal response spectra of S.calcitrans to single and blend odor, the diagrammatic representation of a typical EAG showing parameters used in analysis, amplitude (mV) and response duration in second. (B) Mean EAG amplitudes for the various odor and air (control), n=10. (C). Mean response duration, or time to recovery, n=10 (D). The behavioral response of Stomoxys spp. to single component and blend under field condition, n=4. Error bar represents standard error of the mean.

### Integration of visual and olfactory cues improved trap catch

Once we modified the trap to red, we aimed to enhance the trap catch by baiting it with livestock host odors. In our previous study we identified selective attractants, such as attractant to gravid and blood meal searching *Stomoxys calcitrans.* We used liquid formulation in which 2ml pure odor was loaded every day as a positive control. We formulated a blend of carvone and p-cresol, to target both blood meal searching as well as gravid females for maximum impact. The impregnation process involved the addition of 800 µl of a blend in a 1:1 ratio to 4 grams of nanopolymer beads, following the procedures outlined in the techniques section. As a positive control 2ml of blend liquid formulation was dispensed from 4ml vial with cotton roll stopper and for liquid formulation we reloaded the odor every day, the same as in our previous study that was required for maximum Stomoxys flies catch, replicate was done by days for five days. Traps position was moved every day to minimize position effect. Both the dry formulation and liquid formulation caught significantly more Stomoxys as compared to unbaited control, ANOVA F=17.33, P=0.0002 (Fig.7A.) and house fly (Fig 7B (F=7.03, P=0.007). Furthermore, the utilization of nanopolymer beads improved the attractivity of semiochemicals to Stomoxys by doubling the catch as compared to liquid formulation, even though not statistically significant (Fig 7A-B). The use of nanobeads reduced the amount of odors to be used, as no odor reloading every day. The nanopolymer dispenser also works for other previously identified attractants such as cymene-p, naphthalene, camphene, camphor, α-pinene all performed very well in nanopolymer beads with significant Stomoxys flies catch (data not shown).

**Fig. 7.**
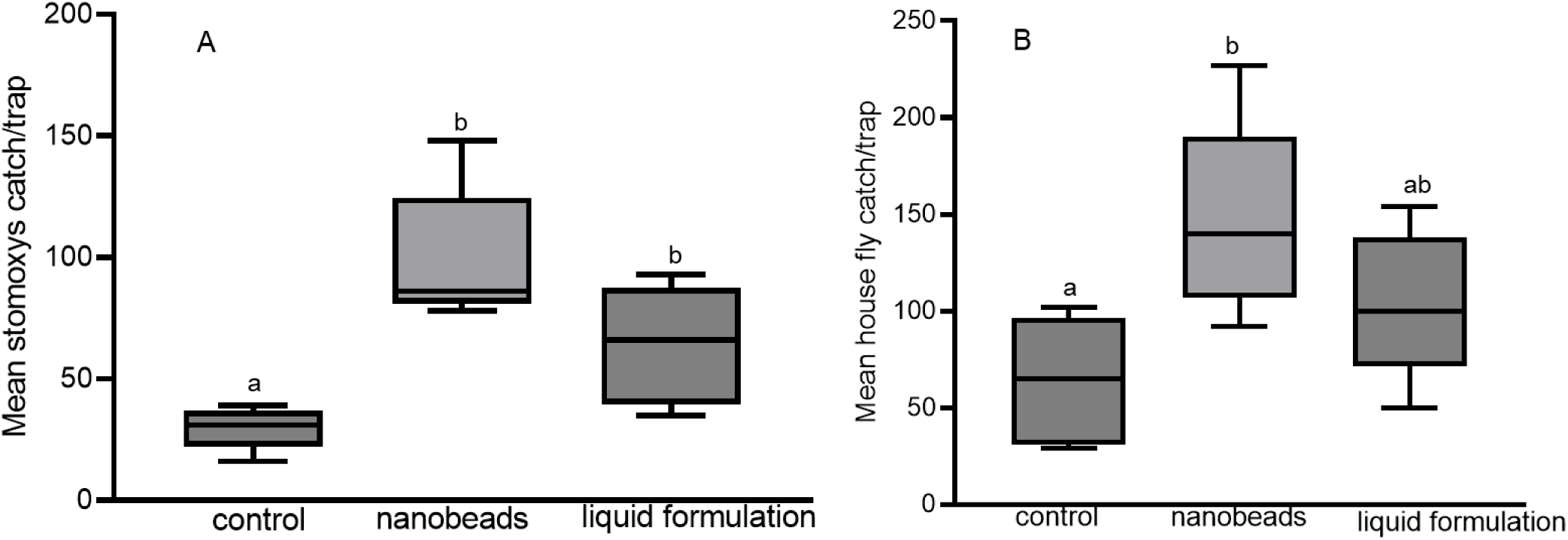
Odors dispensed from nanobeads improved Stomoxys and house flies catch. (A) Response of Stomoxys to red monoconcial trap baited with carvone and p-cresol using two dispensers (nano beads and liquid formulation), at Ngurunit site (B) house flies response.

### The attractivity of red Vavoua trap is independent of ecology

We next challenged our new trap and dispenser at different ecology at two independent sites in Shimba Hills coastal Kenya humid environment as compared to Ngurunit and Nyanuki which are semi-arid ecologies. Red fabric performed more as compared to blue in catching Stomoxys, independent t-test, 2.969, P = 0.01 at Mawia (S: 042100.8, E: 0391820.2) sites and at Tawani site (S: 041742.4, E: 0392647.7) independent t-test 2.986, P=0.009 (Fig 8A). At these two sites located in Shimba Hills similarly the same as Ngurunit site baited traps attracted significantly more Stomoxys as compared to the negative control, F=12.93, P <0.0001 (Fig 8B.). These data demonstrate that the attractivity of red fabric, nanobead dispenser and attractant is independent of ecologies. Unlike the other site red baited trap caught more Stomoxys as compared to blue, t=6.49, P<0.001. The non-target insects caught were very small in both colored traps at Shimba Hill unlike Nanyuki and Ngurunit sites, therefore, no statistical analysis was conducted.

**Figure 8.**
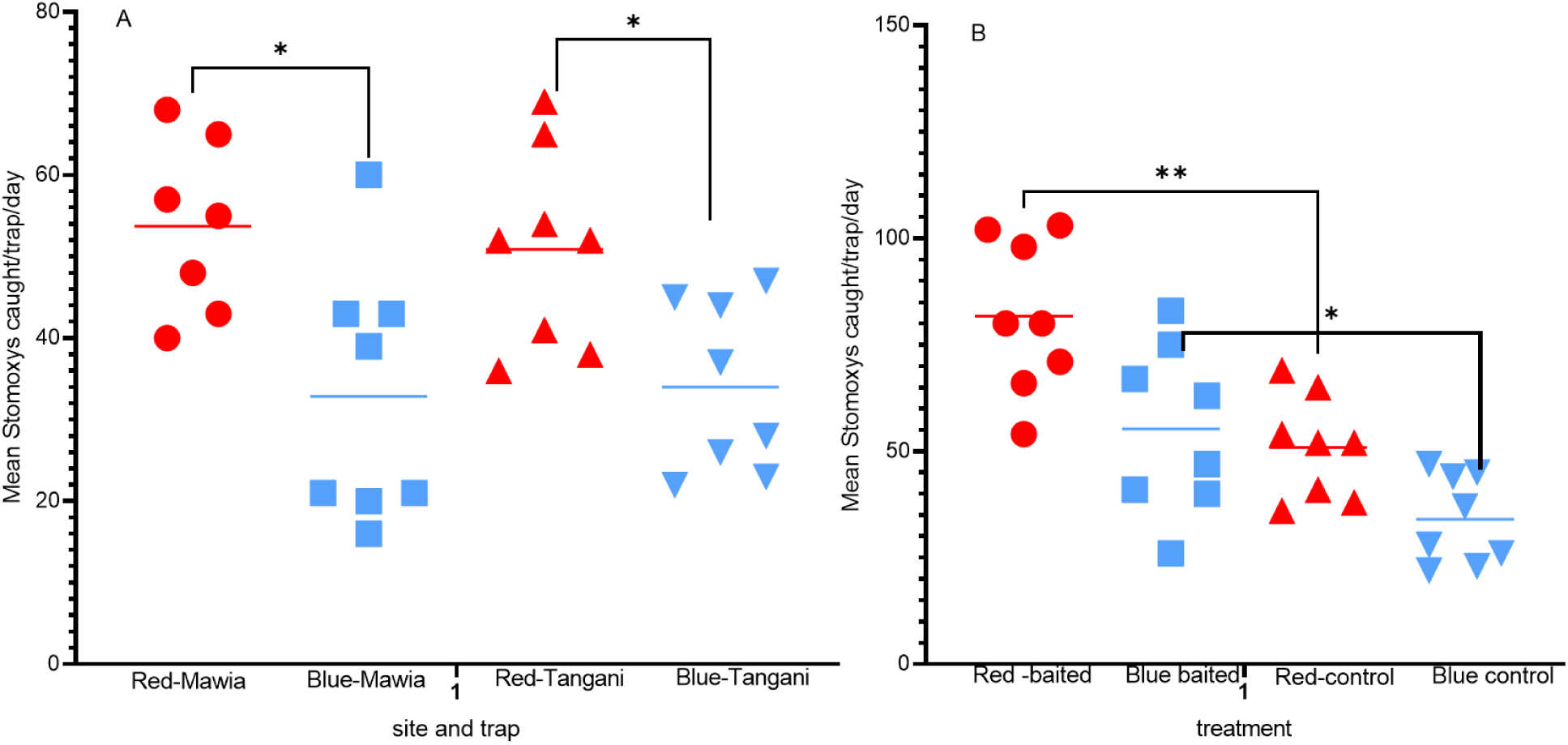
The attractivity of red and blue trap to Stomoxys species at two different sites in coastal Kenya, Shimba Hills. (A) unbaited and (B) baited with blend formulation.

## Discussion

Identifying and locating objects of interest are the most fundamental tasks an insect brain can perform. Insects make use of their various sensory modalities (vision, olfaction, taste etc.) to make measurements and integration of complex and noisy biological signals to solve complex problems to make decision, such as tracking hosts, avoiding enemies, selection of birthing place, and mate partners. Here we report the modification of the Vavoua trap color (the visual cue) from blue to red and dispensing host odors from nanopolymer beads dispenser increased both the efficacy and selectivity of Vavoua trap. Beside livestock and nectar semio-chemicals we hypothesis that the Stomoxys exploit livestock skin color and flower color of nectar sources, which varies in their visible light spectrum is likely important for host recognition and localization at close range, was not supported by our data, as Stomoxys fed equally on various livestock color that various in visual spectrum. The yellow and green fabric that potentially represent some plants flower and leaf color was not attractive to stomoxys. Except black and brown furred cattles the reflectance increased steadily but shows no spectral peaks, the same as [40], that observed similar results in various birds and mammals fur. The observed diversity in cattle fur color can be attributed to variations in pigmentation, specifically melanins, eumelanin and pheomelanin, which all contribute to different fur color phenotypes, in livestock fur [40,41]. However, based on our result it is possible that these color variations have a limited impact on stomoxys - cattle preference visually, as all equally attractive unlike the fabric.

We identified three main species of Stomoxys, the two are the most dominant species in the African continent[24,25,42] however, Stomoxys species composition and density is location, season and methods of collection dependent. The equal attraction of Stomoxys to various livestock fur and four fabric colors that varies in wavelength from (400 – 700nm) may demonstrate that Stomoxys visual system has a broader range of wavelength detection and tuning. Livestock-Stomoxys interaction experiment demonstrated that livestock colors are not essential for the assessment of host but can be attracted to combinations of cues obtained from host such as visual and semio-chemicals. From our previous study we did not find semio-chemical differences between cattle of various colors, such as in their urine, dung, and breath odor profile [21] [43].

However, tsetse flies, which is exclusively a blood-feeder have a differential feeding preference to some animals over others regardless of their abundance [44,45], it seems a combination of visual and olfactory inputs determine the host attractivity, some works demonstrated the impact of odors [20],[46] [47], there is also a strong evidence of visual cue as intensity and angle of polarized lights determine the attractivity of host to biting flies such as tabanus, Stomoxys [48,49]. Furthermore, polarized light combined with size and number of spotty of various color coats, which are widespread among mammals[50] has been shown to determine the attractivity of host to biting flies [51]. Important visual factors are size [52], shape[46], contrast and color [53,54], pattern[55] (Gibson, 1992), and movement[57]. Host defensive behavior has also been speculated to determine the host feeding behaviour of hematophagous insects [6]

Similarly,[12] observed various hematophagous flies attractions to various fabrics of different wavelengths. These common or overlapping perceptual groupings for the four different colors may result from the fact that Stomoxys visual systems exploit similar properties of natural images[58]. But still some fabric that potentially resemble host color such as black, brown, green, yellow did not attract a significant number of Stomoxys, demonstrating beside color there is missing additional features these authors not able to identify, may be such as texture that flies extract to make decision[58]. Further to note in this experiment we did not quantify biting flies that are attracted, but not trapped, as [59] showed it is important to document the fly’s behaviour with video to determine the trap efficiency better. The absence of significant attraction to green and yellow color, which represent some nectar source color indicates flowers may use additional features such as scent to attract pollinators [60–62]. Similarly, [4] showed an apple flies can orient in the absence of visual cues by using only directional airflow cues but require simultaneous odor and directional airflow input for plume following to a host volatile blend. The variation in attraction between various insects to the different fabric colors observed in this study shows insects’ including hematophagous insects’ preference for color and wavelength varies. In tsetse flies slight change in the blue fabric color and associated wavelength accompanied by significant catch difference demonstrating the importance of spectral intensity[10,12,63,64]. This variation in visual cues between insects may be caused by the differing numbers of ommatidia toward the detection of color, and light intensity, which might depend on the spectral sensitivities and interplay of the participating photoreceptors [65–67]. Despite a difference in wavelength, we found red, kaki and blue color were equally attractive to Stomoxys. In agreement to our finding [68]), also reported equal attraction of stable fly to blue and red colored board [69,70] showed red-brown cow was the preferred color for some biting flies. Previously red color was assumed to be invisible to insects, however, recent studies demonstrated that other dipterans, such as model *D. melanogaster* is able to detect wavelengths of red light[71]. In support of our finding electroretinographic recordings from stable flies showed strong peaks of visual sensitivities occurring around 605–635 nm, which is red color zone and at UV zone[72,73].This may necessitate to make some adjustment in our future behavioral experiments that uses red light to simulate darkness[74]. Other researchers also demonstrated the wide color preference of stable flies, for instance[75] demonstrated that white coroplast, and even gray ones, were more attractive to Stomoxys than blue coroplasts. In agreement with this we also show white/gray color is equally attractive to Stomoxys the same as blue, but equally attractive to other non-target insects such as coleoptera.

To attract pollinators via deception principles, plants especially orchids have made various complicated evolutionary adaptation that seems very unlikely, including producing the pheromone of an insect’s shape of a female insects to attract male for mating[76] for review), even heat of dead carcass[14], but they do not reward for the service rendered. However, less complicated objects such as traps and target with the same false signals of reward, that do not look like or smell the blood or nectar source to deceive vectors (Stomoxys, tsetse flies), showing the variation between insects to be deceived. We have observed a synergism effect due to the blend as compared to single component under field condition in behavioral response, however, the EAG response of the blend did not change from single component, this may be athough EAGs show a concentration– response relationship with stimulus concentration[36], the EAG response represent qualitative, rather than quantitative indicator of olfactory response.

Insect vectors navigation to their host and traps is affected by upwind flight due to the intensity of molecular flux of individual odor strands[77][78], that need focusing on producing dispensers that would create the strongest possible strands downwind for maximum behavioral impact. The use of nanopolymer beads as demonstrated by the behavioral efficacy, odor integrity and longevity may result in an increase odor strand, in dispensers’ field longevity while reducing the quantity of expensive odors that is used per dispenser. The improved attraction of the same odor when used nanobeads as compared to the other two dispensers, may be accounted due to the small size of the nanobeads as compared to both wax and cotton roll, which will create small point source for the odors, and maintain high release of the odor with strong strand that has more behavioural impacts[34]. The geometries of dispensers and their alignment with respect to the wind line may be another way to optimize dispensers’ abilities to create strong plume strands and thereby potentially use the semio-chemicals in the dispensers more efficiently[78] [77][34]. In our trial the use of circular tygon tube to dispense the attractant odors from nanobeads may be another addition in optimizing emission rates and efficacy allowed maximizing dispenser exposure to the environment such as directional airflow, which is required for plume following. Here we show nanobead polymer as a potential dispenser because of its slow release of the target odor(s), with a strong odor plume, which keeps the odor integrity and minimize cost.

## Conclusions

A significant attraction of *Stomoxys spp*. to various colors, but red demonstrated high efficacy and selectivity to *Stomoxys spp*., independent of ecologies demonstrating it is an environmentally preferred trap and may be used to combat vector borne diseases such as animal trypanosomiasis[79]. Host odor blend dispensed from nanopolymer bead significantly increased trap catch, demonstrating the importance of integrating multimodal signals (odor and visual) for maximum *Stomoxys* attraction. Interestingly, *Stomoxys* were avoiding some colors, showing the *Stomoxys* visual system is a promising target for selective attraction and inhibiting their attraction to animal hosts or animals’ enclosure for instance in zero grazing system. Furthermore, we demonstrated nanobeads as economical dispenser with high efficacy and suitability for field application in an economical way.

## Ethical clearance

The animal study was reviewed and approved by Animal Care and Use Committee (IACUC) of the International Centre of Insect Physiology and Ecology, reference: IcipeACUC2018-003-2023. Verbal informed consent was obtained from the owners for the participation of their animals in this study.

## Supporting information

Suplimentary table1 livetsock Refelectance data .rar

Suplemntary table 2 Stomoxys color attarction Field work data.rar

## Data accessibility

All data are included in the manuscript and in Supplementary materials.

## Declaration of AI use

We have not used AI-assisted technologies in creating this article.

### Author contribution

MNG: Conceptualized, designed, experimented, analyzed, wrote the manuscript, and fund mobilization. SBB, designed, conducted fieldwork, and analyzed data. JN, PA contributed in field work. DM designed and fund mobilization. All authors read and commented on the manuscript.

## Acknowledgment

We thank Dr Steve Mihok for his critical and significant review, comments, and suggestions on the original draft. We thank Emily, Drs Bestel and Elfathi for their useful discussion about getting spectrum measurement. We thank Victor Omondi, JohnMark and Simon Tawich for their useful technical help in getting spectrum measurement from livestock and fabric.

