## Supplementary table 3 for "Multimodal interactions in Stomoxys navigation reveals synergy between olfaction and vision"

Supplementary Table 3. Electrophysiological response of *S.calcitrans* to host odours

| compound | Response(mV) | Response duration(sec) |
|---|---|---|
| control | 0.31 | 0.47 |
| control | 0.11 | 0.89 |
| control | 0.29 | 0.35 |
| control | 0.1 | 0.35 |
| control | 0.09 | 0.7 |
| control | 0.41 | 0.55 |
| control | 0.22 | 0.58 |
| p-cresol | 1.47 | 2.82 |
| p-cresol | 1.36 | 3.06 |
| p-cresol | 1.46 | 4.12 |
| p-cresol | 1.19 | 4.57 |
| p-cresol | 1.58 | 2.64 |
| p-cresol | 0.37 | 1.4 |
| p-cresol | 2.29 | 3.53 |
| p-cresol | 2.6 | 3.87 |
| p-cresol | 1.28 | 2.64 |
| p-cresol | 2.34 | 3.6 |
| carvone | 1.78 | 4.16 |
| carvone | 1.99 | 5 |
| carvone | 1.55 | 4.3 |
| carvone | 1.69 | 3.64 |
| carvone | 1.49 | 4.47 |
| carvone | 0.94 | 2.3 |
| carvone | 4.3 | 4.52 |
| carvone | 3.65 | 5.61 |
| carvone | 1.31 | 2.82 |
| carvone | 3.69 | 3.7 |
| New blend | 1.28 | 2.47 |
| New blend | 1.55 | 3.19 |
| New blend | 1.45 | 2.59 |
| New blend | 1.42 | 2.82 |
| New blend | 1.17 | 2.3 |
| New blend | 0.72 | 2.37 |
| New blend | 3.95 | 3.7 |
| New blend | 3.44 | 3.41 |
| New blend | 1.34 | 3.35 |
| New blend | 3.65 | 3.06 |
| usedblend | 1.68 | 3.04 |
| usedblend | 1.7 | 2.35 |
| usedblend | 1.48 | 2.24 |
| usedblend | 1.75 | 3.19 |
| usedblend | 1.18 | 1.94 |
| usedblend | 0.77 | 2.43 |
| usedblend | 4.18 | 3.17 |
| usedblend | 3.68 | 2.47 |

| | | |
|---|---|---|
| usedblend | 1.15 | 3.3 |
| usedblend | 3.8 | 3.06 |
